# Optimizing Consensus Generation Algorithms for Highly Variable Amino Acid Sequence Clusters

**DOI:** 10.1101/2020.11.08.373092

**Authors:** Reyhaneh Mohabati, Reza Rezaei, Nasir Mohajel, Mohammad Mehdi Ranjbar, Kayhan Azadmanesh, Farzin Roohvand

## Abstract

Producing a functional consensus sequence is a preliminary bioinformatics task, which is a necessity for many research purposes. However, the existence of hypervariable regions in the input multiple sequence alignment files causes complications in generating a useful consensus sequence. The current methods for consensus generation, Threshold, and majority algorithms, have several problems, which exclude them as applicable algorithms for such highly variable sequence clusters. Hence, we designed a novel alternative algorithm for the same purpose. The algorithm was explained both using a mathematical formula and a practical implementation in Python programming language. A sequence set from HCV genotype 1b E2 protein has been utilized as a practical example for evaluating the algorithm’s performance. A few in silico tests have been performed on the output sequence and the results have been compared to results from other algorithms. Epitope-mapping analysis indicates the functionality of this algorithm, by preserving the hotspot residues in the consensus sequence, and the antigenicity index shows significant antigenicity of the consensus sequence. Moreover, phylogenetic analysis shows no significant change in the placement of the new consensus sequence on the phylogenetic tree compared to other algorithms. This approach will have several implications in designing a new vaccine for highly variable viruses such as HIV-1, Influenza, and Hepatitis C Viruses (HCV).

## Introduction

Finding an average sequence, consensus, by calculating the most frequent residues in each position, both for nucleic acids and polypeptide sequences, is a common bioinformatics task(1). Finding common structural motifs between DNA and protein sequences(2,3), designing a set of primers, which will attach to a set of nucleotide sequences(4), or determining the hotspots for antibody-epitope interactions in a set of sequences related to a pathogen are just some of the applications of consensus sequences. But the major reason for generating a consensus sequence is minimizing the genetic distances of highly variable or hypervariable regions of a nucleotide or polypeptide sequence, which have the most amount of heterogeneity among related sequences(5–9). In many cases, the variability of these regions is an evolutionary advantage for the sequence containing them. However, if that sequence is in the immunodominant proteins of a pathogen, it will be a major obstacle for designing effective broad coverage vaccines against most strains of that pathogen. So, consensus sequences are utilized to overcome the genetic diversity of variable sequences. To this end, several different bioinformatics software packages were designed to generate a consensus sequence from a set of input sequences. Most of these software packages use either Majority or Threshold-based selection methods(10–12) as their underlying algorithm. In the threshold-based method, the algorithm chooses the residue, which has a higher frequency than the user’s selected threshold while the Majority selection method chooses the most common residue in each position regardless of any indicated threshold. Even though these two algorithms have been effectively used in a wide variety of applications, their selection accuracy for highly variable sequence clusters is controversial. For example, the rigidity of the threshold-based selection algorithm on a specific threshold point might lead to neglecting some residues with very close frequencies, but slightly lower than the selected threshold. Moreover, consensus sequences generated through a threshold-based algorithm might have no functional use because of the gaps created in residues with the frequencies lower than the selected threshold. Majority based selection has also some disadvantages such as selecting exclusively based on the frequency differences without considering the population weight and producing a polypeptide with highly different conformational and functional forms.

So, there is a need to design a new algorithm that resolves the problems of the threshold-based method by having the flexibility of the Majority selection algorithm and gives reliability to the selection process by involving the natural selection and natural priority of each residue in a given position. The algorithm gives a quantitative measure of how fit an amino acid residue is for its corresponding position. Moreover, a variable sequence cluster from glycoprotein E2 of the Hepatitis C Virus, protein sequences of subtype 1b from genotype 1, was considered to evaluate the accuracy of the new algorithm. A comparison of the new algorithm to previous consensus algorithms has been studied, too.

## Methods

### 1 The Fitness Score

The basis of this new algorithm is calculating a fitness score for each residue in a position and selecting the fittest residue for that position. The procedure will be explained for one position and it can be reiterated for the rest of the positions for any sequence length.

The fitness score is calculated by considering each residue as the possible candidate for that position and then calculating the tendency of natural selection to keep that amino acid in that position. This tendency is a function of residue’s frequency and its substitution score, which will be obtained from BLOSUMs (Blocks Substitution Matrices). BLOSUMs, amino acid substitution matrices, contain every possible amino acid pairs with a quantitative measure of their substitution likelihood. Moreover, based on the evolutionary distance of the input sequences, the most suitable matrix will be chosen, for example, BLOSUM 30 for very far sequences and BLOSUM 80 for very close ones. The algorithm multiplies each possible substitution pair score, including the substitution of amino acid with itself, with the frequency of the base amino acid. Then, the total fitness score will be calculated by adding each residue’s corresponding scores. Finally, the residue with the highest fitness score will be chosen.

The process for one imaginary position would be as follows:

1. BLOSUM62 matrix will be chosen for convenience but BLOSUM 30, 45, 60, and 80 can be used.
2. This position has four different amino acid residues in different sequences, including ‘G’, ‘W’, ‘T’, and ‘A’.
3. The frequency of each residue is: ‘G’: 0.142, ‘W’: 0.142, ‘T’: 0.428, ‘A’:0.285
4. A table will be created using the letter names both as row names and as column names. By doing so, each cell of this table will be a possible combination pair including self-pairs (Fig1-A).
5. The substitution score of each pair will be retrieved from the BLOSUM62 matrix and placed in its corresponding cell(s), one cell for self-pairs in the diagonal, and two cells for other pairs (Fig1-B).
6. The frequency of each row’s residue will be multiplied in the corresponding substitution scores in that row. For example, the 0.14 frequency of G residue will be multiplied in each cell in that row (Fig1-C).
7. Calculate the total fitness score for each amino acid in each column by adding the corresponding cells of that column together (Fig1-D).
8. The residue with the highest fitness score will be chosen as the representative of that position in the final consensus sequence.

**Figure 1.**
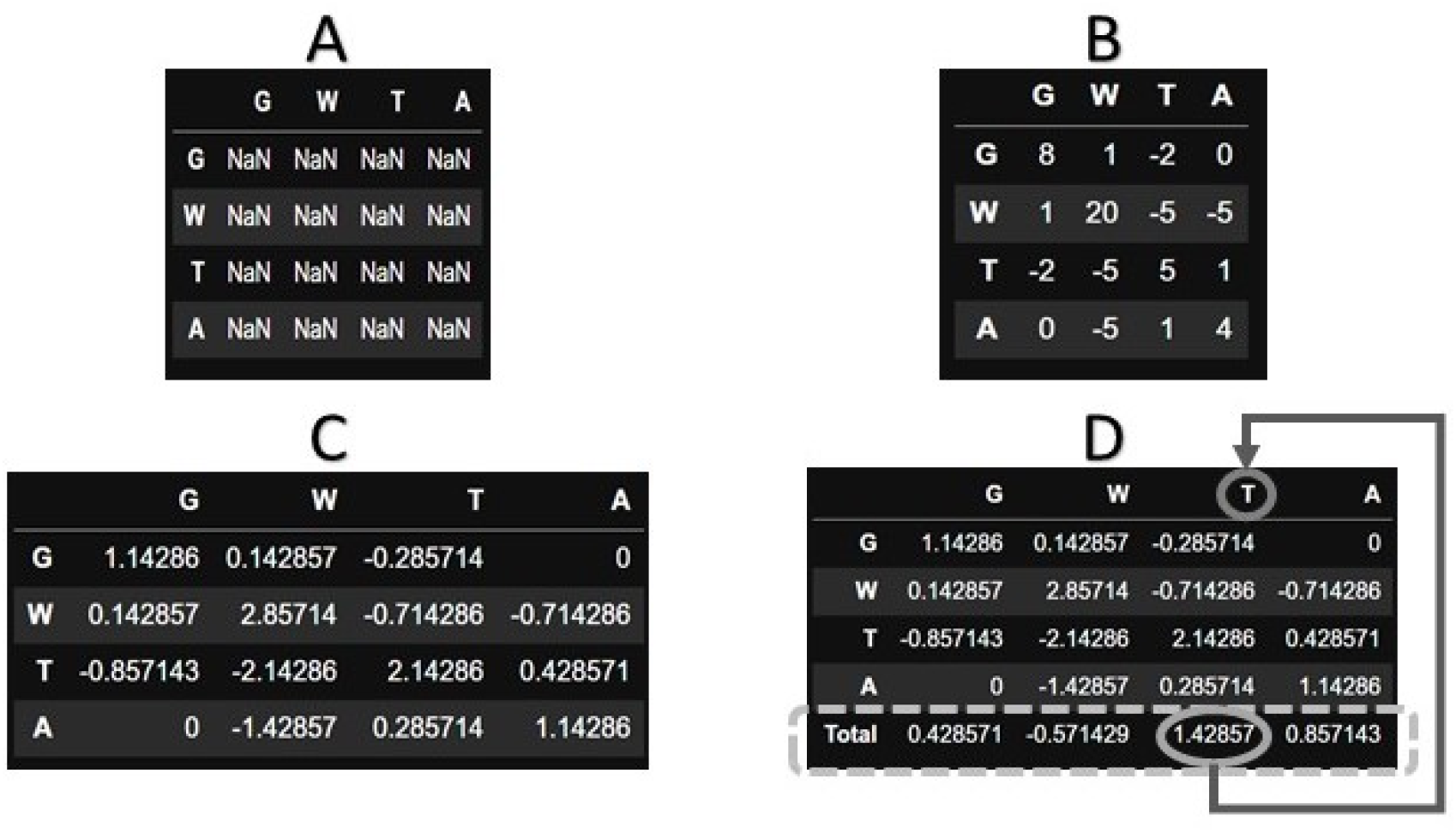
A: the empty table - B: each cell contains the corresponding substitution score of its row and column pair - C: each cell in a row has been multiplied with the frequency of its corresponding row name - D: the last column contains the total of each column, the max fitness score and its corresponding column name has been highlighted.

The above procedure was a detailed explanation for implementing this algorithm in a programming language. However, the fitness score for each residue can be calculated by the following formula:

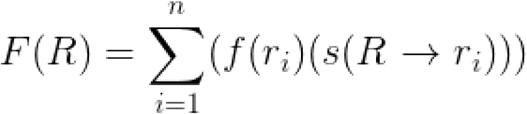

*Equation 1. F: fitness score function, R: target residue, f: frequency function, r: each residue including the target itself, R −> r: substitution of the ‘R’, or target, residue with the ‘r’ residue.*

#### Consensus Generation Pipeline

The basic assumption of the fitness algorithm is getting an aligned sequence set, multiple sequence alignment (MSA) file, as its input. The MSA file should be in a familiar format to let the algorithm extract the necessary information, including sequence length, frequency of each residue in any given position and the list of existing residues in each position. Therefore, to make a consistent pipeline, the alignment part is also added to the base algorithm to make it more robust and reliable. The MAFFT alignment python package was used as the basic alignment tool because it was the most flexible alignment algorithm with several changeable options, which could be effective in the result of the fitness algorithm function. One of the most important options of the MAFFT algorithm was the ability to change the substitution matrix, which is being used in aligning the given sequence set. The substitution matrices in the MAFFT algorithm, which can be chosen based on the evolutionary distance between the input sequences, includes BLOSUM30, BLOSUM45, BLOSUM 62, BLOSUM80, and other scoring matrices. The chosen matrix can be the same or different from the one which is being used in the fitness calculation process, which will add extra flexibility to the final pipeline.

The consensus generation pipeline including the preprocessing steps, the score calculation steps, and the consensus sequence export has been implemented in python programming language. All necessary steps in the pipeline are summarized in the flowchart format (Figure 2).

**Figure 2.**
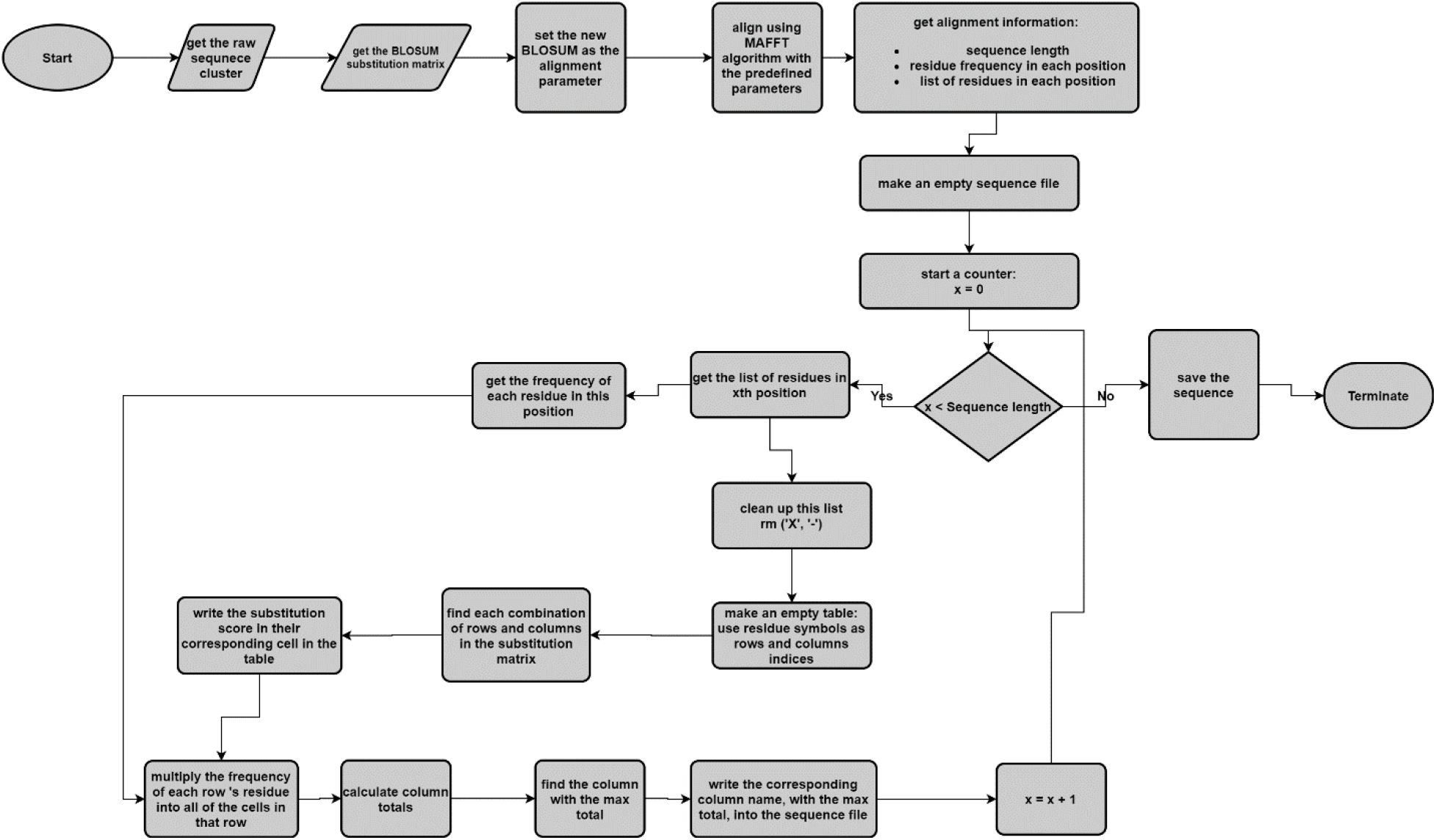
The implementation of the consensus generation procedure has been depicted in the flowchart format.

### 2 Generation of consensus sequences for Hepatitis C virus (HCV) E2-1b cluster

More than 400 HCV-E2-G1b sequences were retrieved from Virus Pathogen Database and Analysis Resource (ViPR)^1^. The retrieved sequence set was aligned using the MAFFT algorithm^2^ (13)and went through the consensus generation process using three algorithms. CLC genomic workbench 5.5^3^ was utilized to generate six consensus sequences based on the threshold (50-100%) algorithm and one consensus sequence based on Majority algorithm. The fitness algorithm was also separately applied to generate a new consensus sequence.

### 3 Phylogenetic Tree Analysis

Applying iqtree^4^ software(14), the phylogenetic distance of consensus sequences were calculated with both SH-aLRT and UFBoot support values. The phylogenetic tree was rendered using FigTree software^5^.

### 4 Receptor and Antibody Binding Important Residues

The necessary residues for antigen-antibody interaction and also the residues involving in receptor binding sites were considered to evaluate the accuracy of the algorithm in preserving the important residues. To this end, one neutralizing antibody named (AP33) (15,16) targeting the linear epitope and two neutralizing antibodies named (1:7), which target the same residues that are also necessary for CD81 receptor binding, (17–19) and (HC-84.1)(20) targeting conformational epitopes were studied.

### 5 Antigenicity prediction

The potency of consensus sequence to be an antigen was tested through Vaxijen (Prediction of Protective Antigens and Subunit Vaccines)^6^ and AntigenPro^7^ servers.

### 6 Prediction of N-glycosylation sites in E2 proteins

To determine their association with viral neutralization, the N-glycosylation sites in E2 protein were predicted using NetNGlyc Server 1.0^8^.

## Results

### 1 Alignment of consensus sequences

The aligned representation of amino acids of all consensus sequences was illustrated in Figure 3. The first impression of this illustration shows the weakness and inapplicability of the threshold algorithm. The number of undefined amino acids in threshold-based consensus sequences is too many to make any applicable sequence, which could lead to a functional epitope. Even though both the Majority algorithm and the fitness algorithm, named G1b Blusom62 in figure 3, produce operative sequences, their outputs are significantly different which is an indication of the different consensus generation approach in the fitness algorithm.

**Figure 3.**
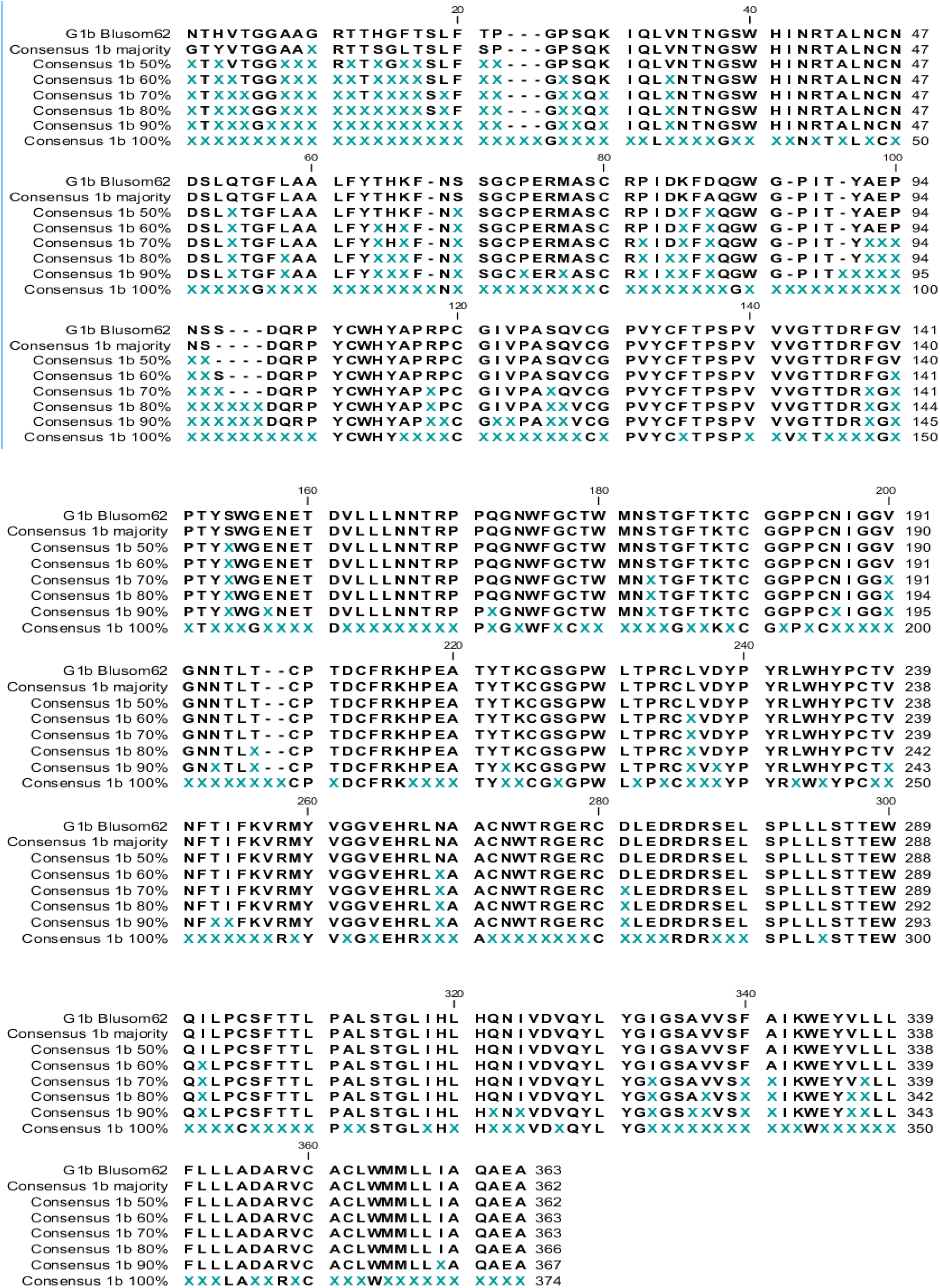
Comparing consensus sequences generated from more than 400 HCV-E2-G1b sequences through different algorithms. X: undefined amino acid, Blosum based Con-Seq: Fitness algorithm, Majority based Con-Seq: Majority Algorithm, Threshold-based Con-Seq: Threshold Algorithm.

**Figure 4.**
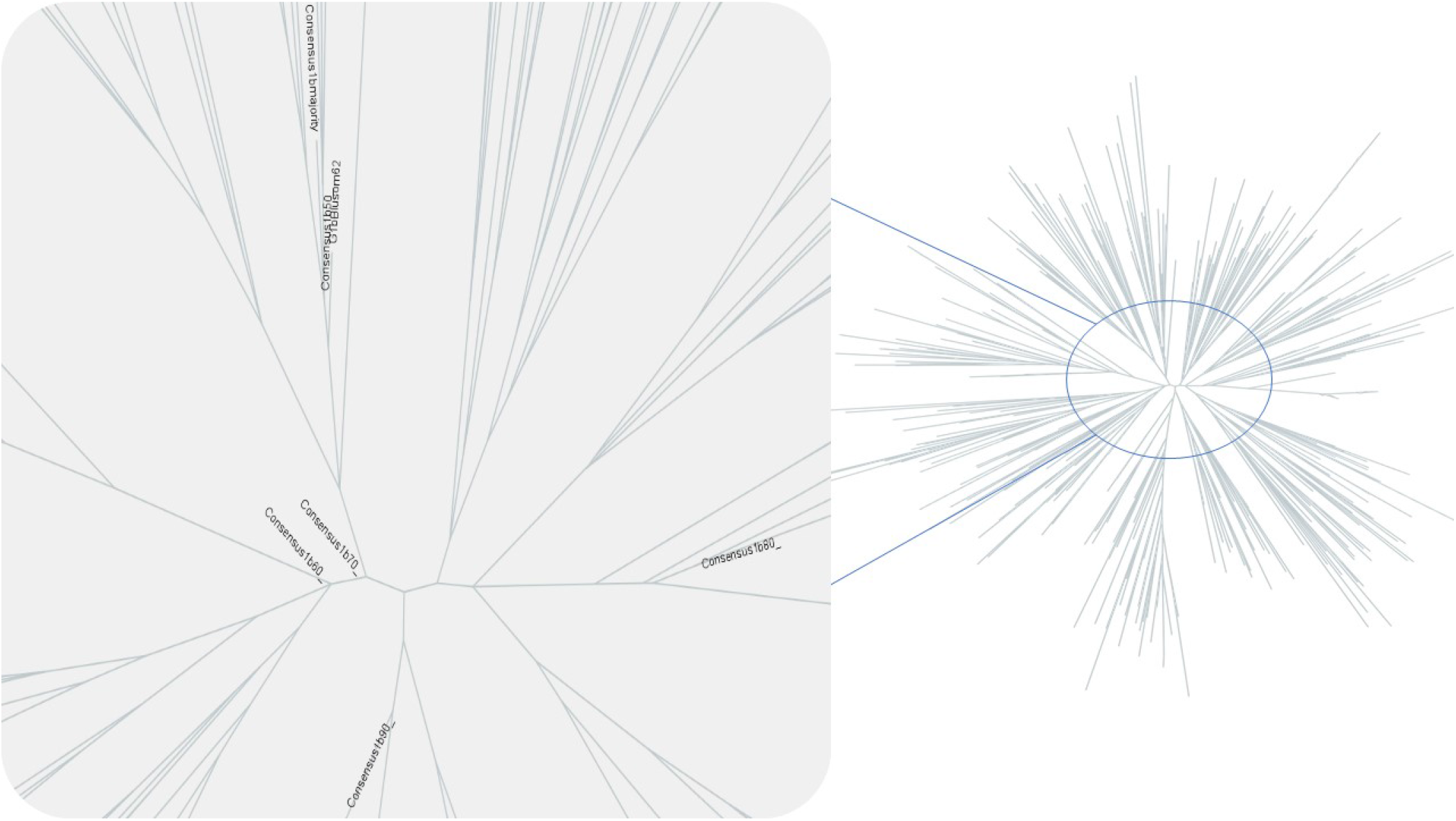
Phylogenetic Tree. The unrooted phylogenetic tree was inferred from E2 amino acid sequences and the consensus sequences generated based on three algorithms. Sequences were aligned with MAFFT algorithm and iqtree has been used for phylogenetic tree construction.

Figure 3 shows a qualitative comparison between the fitness algorithm and other algorithms. Although it is impossible to determine the efficiency of an algorithm without utilizing its output in several downstream procedures, in vitro, and in vivo tests, the fact that its result shows significant differences compared to the previous algorithms could be persuasive enough for further investigations.

### 2 Comparison of Phylogenetic Trees

The distance between consensus sequences generated through different algorithms on the Phylogenetic tree seems negligible. This result implies that the fitness algorithm does not significantly change the location of the consensus on the tree even though it generates a different sequence. The 100% Threshold consensus sequence contains too many gaps and it failed the phylogenetic distance calculation in iqtree software and therefore it is not presented in the phylogenetic tree.

### 3 Conserved residues Preservation Potency

As illustrated in Figure 5, no change occurred in important residues responsible for interaction between antigen and three selected neutralizing antibodies in this study (Figure 5). Normally, these residues are conserved in the protein sequence and this result shows that the fitness algorithms preserve the sequence of conserved regions.

**Figure 5.**
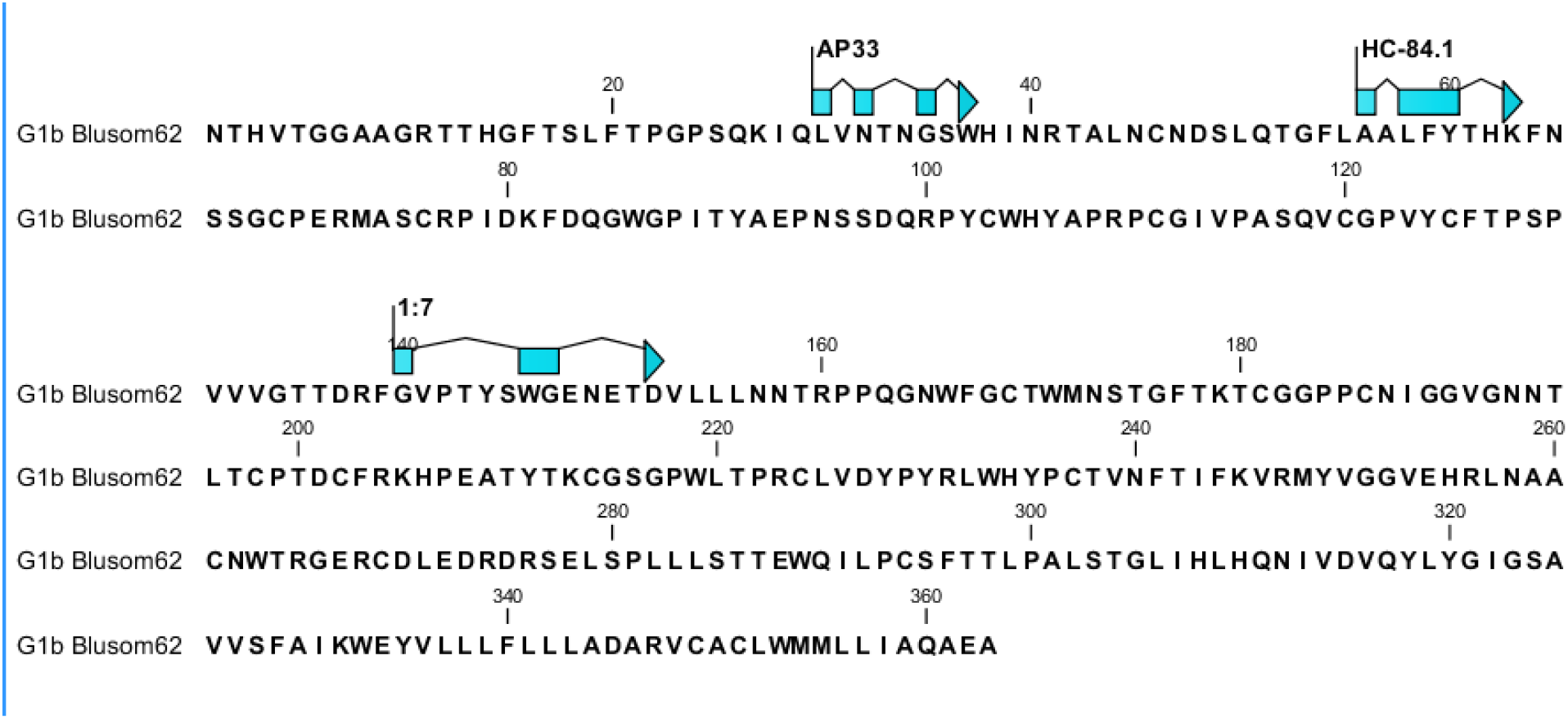
Conserved Residues – Arrows indicate the residues involved in epitope-antibody interaction.

### 4 Antigenicity Evaluation

Data obtained from online servers (Vaxijen and AntigenPro) were indicated that the antigenicity index of consensus sequence is above the thresholds considered by the servers (table1). This result also shows that the fitness algorithm has very close antigenicity score to G1b Threshold 90% in both of these tests.

**Table 1.** Antigenicity of all consensus sequences predicted by Vaxijen and AntigenPro. Vaxijen threshold is 0.4 and AntigenPro threshold is 0.5. *: Non-Antigen *: Consensus Sequences generated from three algorithms.

| Sequences*<br>Servers | G1b<br>Blosum<br>62 | G1b<br>Majority | G1b<br>Threshold<br>d 50% | G1b<br>Threshold<br>d60% | G1b<br>Threshold<br>70% | G1b<br>Threshold<br>80% | G1b<br>Threshold<br>90% | G1b<br>Threshold<br>100% |
| --- | --- | --- | --- | --- | --- | --- | --- | --- |
| VaxiJen | 0.5208 | 0.4934 | 0.5760 | 0.5848 | 0.5936 | 0.5813 | 0.5411 | 0.3597* |
| AntigenPro | 0.7867 | 0.7625 | 0.6640 | 0.7224 | 0.7687 | 0.7430 | 0.8016 | 0.7773 |

### 5 Predicted N-glycosylation Sites

NetNGlyc server was applied to predict the sites under N-glycosylation modifications. The number of glycosylation sites on the E2 proteins are between 9-11 sites. All algorithms, except Threshold 100%, preserve at least 9 sites. However, the fitness algorithm preserves all of the 11 reported glycosylation sites including the E2N5 position which has higher variability in 1b sequences(21). Therefore, the fitness algorithm result saves the most possible natural glycosylation sites of this protein. (Figure 6)

**Figure 6:**
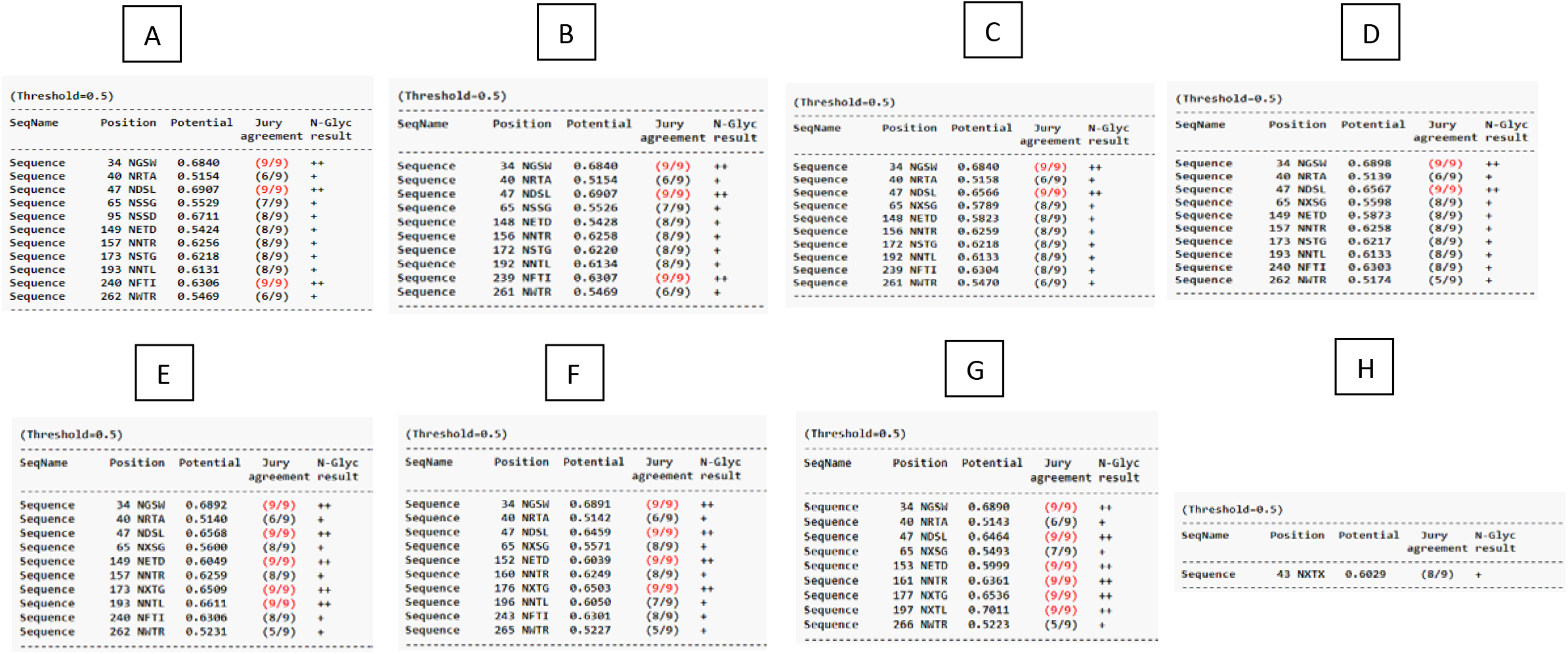
The sites of N-glycosylation were shown in all of the consensus sequences generated from three algorithms. **A**: G1b Blosum62. **B**: G1b Majority. **C**: G1b Threshold 50%. **D**: G1b Threshold60%. **E**: G1b Threshold 70%. **F**: G1b Threshold 80%. **G**: G1b Threshold 90%. **H**: G1b Threshold 100%.

## Discussion

In current research, a new innovative consensus generation algorithm, fitness algorithm, was introduced, and its efficacy and differences from the previous methods were demonstrated, using variable HCV E2 subtype 1b sequences as an example.

generating an intermediate canonical sequence, which preserves the features of the original variable regions is a challenge that the current consensus generation algorithms, Threshold and Majority, haven’t been able to solve.

The disadvantages of the threshold-based selection for such collection of diverse sequences are as follows:

1. In most of the positions, there is no residue with an upper than 60% frequency, which could be chosen by specifying a reliable threshold point, usually 60% or more.
2. The rigidity of this algorithm on a specific threshold point might lead to neglecting some residues, which have frequencies very close, but lower than the selected threshold, for instance a residue with 59.9% frequency will be omitted by a 60% threshold selection.
3. The result consensus sequence will have many gap positions and consequently information loss, which makes this consensus sequence nonfunctional for practical use. Although the majority selection algorithm does not have the mentioned limitations, it does not consider the evolutionary cost of substituting a specific residue with the other candidate residues in a position and only cares about pure proportion calculating the sample population. In the case of two residues with very close frequencies, choosing between them just based on their frequency difference could not be a reliable approach because the frequencies in a set of sample sequences may not accurately represent the actual frequencies in the population. For example, considering two residues one with 48% frequency and the other with 47.5% frequency, the majority algorithm will choose the former even if the real proportion of the latter in their corresponding natural population is significantly more.

Therefore, we attempted to design a more reliable and optimized consensus generation algorithm for highly variable sequences in this research. This new algorithm has the following features:

1. The selection process is flexible enough to fill all o positions, preventing undecided or gap positions and nonfunctional sequence products, and solves the problem of having one static threshold point.
2. The residues are selected not solely based on the frequencies in this collection of sequences but also by considering the relative substitution probability of that residue in nature.

Even though the main aim of the study was to introduce this new algorithm using common mathematical notions, a script was also written to represent a functional version of our algorithm for comparison reasons.

The hepatitis C virus (HCV) is a major global health burden that infects 1–2% of the world population with an estimated 1.5–2 million new infections each year(22–27). HCV vaccine development is challenging due to the high level of diversity caused by an RNA-dependent RNA polymerase (RdRP) that lacks proofreading activity, thus increase the rate of replication errors and generating significant genetic diversity(28–32). consequently, due to the diversity, there are seven genotypes and 67 subtypes that have at least 32.39% and 14.55% nucleotide variation and 25.02% and 9.7% amino acid variation, respectively(33). Moreover, the degree of genetic variation in different regions of the HCV genome varies. The highest degree of genetic heterogeneity is due to the structural gene encoding the envelope glycoprotein 2 of HCV, named E2 (34,35). Major neutralizing epitopes are located in E2 hypervariable regions (HVRs) where only 37% of the positions shared the same consensus amino acid across all HCV genotypes and can rapidly adapt to host immunity, leading to escape(36) or exhibit structural variability(37–39). Of note, antibodies against these regions are strain-specific but a vaccine must elicit antibodies able to neutralize the vast majority of circulating strains (33). Among the HCV subtypes, studies revealed that subtype 1b is highly variable. (40a)(44) Therefore, generating a consensus sequence from subtype 1b, not only decreases the genetic distances among all subtype 1b sequences but also it will take us one step further to generate a vaccine candidate to induce broadly cross-neutralizing antibodies which is essential for a prophylactic vaccine against HCV. Hence, the HCV E2 1b genotypes were used as an example to compare the new fitness algorithm to the other conventional consensus generation algorithms and to give a view on the application of this algorithm for vaccine development studies.

The functional evaluation of the fitness algorithm was done through a set of *In-silico* tests such as epitope mapping, glycosylation, phylogenetic tree, and also antigenicity prediction.

In detail, the new consensus sequence has kept the hot residues necessary for antigen-antibody interactions, whether linear or conformational ones.

The phylogenetic study shows that despite the significant difference in the polypeptide sequence between the new algorithm and the older methods, the overall position of the consensus sequence on the tree is preserved and it is very close to the center of the unrooted tree.

The probability of consensus sequence for being antigen was also determined through two different servers and results indicated that the consensus sequence generated from the fitness algorithm has an antigenicity score above both servers’ thresholds and also near to threshold 90% while the consensus sequence generated from fitness algorithm has no gaps compared to threshold 90% generated consensus sequence.

The glycosylation was also evaluated on the new sequence and the result implies that all reported glycosylation sites have been preserved through the fitness algorithm while other methods show a fewer number of glycosylation sites. Hence, the data indicates a closer correlation between the fitness algorithm and the natural strains compared to the other algorithms.

Altogether, we designed a consensus generation pipeline, named fitness algorithm, and evaluated the generated sequence’s efficacy compared with the one majority-based consensus sequence and six threshold-based consensus sequences via a variable sequences cluster from E2 protein of HCV-subtype 1b(G1b) through a set of in-silico tests.

To use this novel consensus generation algorithm for practical examples, such as vaccine development studies, in vitro, and in vivo studies on the sequences generated from this algorithm are of crucial importance. Hence, the future perspective for this study is to consider the functional difference and applicability of the fitness consensus sequences in the production of cross-protective neutralizing antibodies in model organisms.

### Supplemented Material

The python implementation has been saved in bloConGen.py, which is supplied to this article. The file can be executed or modified using any python terminal or IDE, provided having the following packages installed in the python environment: Biopython, Pandas, and Mafft Commandline.

## Footnotes

1 https://www.viprbrc.org/brc/home.spg?decorator=vipr

2 https://mafft.cbrc.jp/alignment/software/

3 www.clcbio.com

4 http://www.iqtree.org/

5 http://tree.bio.ed.ac.uk/software/figtree/

6 http://www.ddg-pharmfac.net/vaxijen/VaxiJen/VaxiJen.html

7 http://scratch.proteomics.ics.uci.edu/explanation.html~ANTIGENpro

8 http://www.cbs.dtu.dk/services/NetNGlyc/

